# Lingering neural representations of past task features adversely affect future behavior

**DOI:** 10.1101/2022.03.08.483546

**Authors:** Benjamin O. Rangel, Eliot Hazeltine, Jan R. Wessel

**Affiliations:** Interdisciplinary Graduate Program in Neuroscience, University of Iowa, Iowa City, IA 52245; Cognitive Control Collaborative, University of Iowa, Iowa City, IA 52245; Department of Psychological and Brain Sciences, University of Iowa, Iowa City, IA 52245; Department of Neurology, University of Iowa Hospitals and Clinics, Iowa City, IA 52242

## Abstract

During goal-directed behavior, humans purportedly form and retrieve so called ‘event files’ – conjunctive representations that link context-specific information about stimuli, their associated actions, and the expected action-outcomes. The automatic formation – and later retrieval – of such conjunctive ‘event file’ representations can substantially facilitate efficient action selection. However, recent behavioral work suggests that these event-files may also adversely affect future behavior, especially when action requirements have changed between successive instances of the same task context (e.g., during task-switching). Here, we directly tested this hypothesis through a recently developed method that allows measuring the strength of the neural representations of context-specific stimulus-action conjunctions (i.e., event files). Thirty-five male and female adult humans performed a task-switching paradigm while undergoing EEG recordings. Replicating previous behavioral work, we found that changes in action requirements between two spaced repetitions of the same task incurred a significant reaction time cost. By combining multi-variate pattern analysis and representational similarity analysis of the EEG recordings with linear mixed-effects modeling of trial-to-trial behavior, we then found that the magnitude of this behavioral cost was directly proportional to the strength of the conjunctive representation formed during the most recent previous exposure to the same task – i.e., the most recent ‘event file’. This confirms that the formation of conjunctive representations of specific task contexts, stimuli, and actions in the brain can indeed adversely affect future behavior. Moreover, these findings demonstrate the potential of neural decoding of complex task set representations towards the prediction of behavior beyond the current trial.

## Introduction

According to influential theories of human cognition (e.g., Hommel et al., 2001, Frings et al., 2020), humans automatically store and retrieve episodic representations of their interactions with the world to guide future behavior. These representations contain context-specific combinations of stimuli, actions, and outcomes, and are also known as ‘event-files’ (Hommel 2009; Hommel et al. 2001). For instance, in the example of opening a safe, an event-file would contain a conjunction of the specific context (e.g., the room with the safe), a stimulus (the combination lock), an action (entering the combination), and an outcome (the opening of the safe). The formation and retrieval of such conjunctive event-files can greatly facilitate behavior by making repeated actions more efficient.

However, it is also observed that these event-files can have adverse effects on behavior (Frings et al. 2020; Hommel 2004; Hommel and Colzato 2004). For example, when the correct response or its outcome contingencies change between successive exposures to the same context (e.g., if the safe’s combination has been recently changed), preexisting event-files may contain outdated representations and prime an inappropriate action (Hommel et al. 2014; Neill 1997; Waszak et al. 2003). In the laboratory, these effects can be measured in the N-2 Repetition Cost (N2RC) paradigm (Grange et al. 2013; Kowalczyk and Grange 2016; Mayr 2002; Mayr and Keele 2000). In such experiments, triplets of successive trials made in separate task contexts (C-B-A) are compared with triplets of trials that include a repetition of task A (A-B-A). The N2RC effect is a reaction time cost on the third trial of an ABA compared to a CBA triplet when the response on third trial differs from the response on the first trial (***Figure 1***).

**Figure 1.**
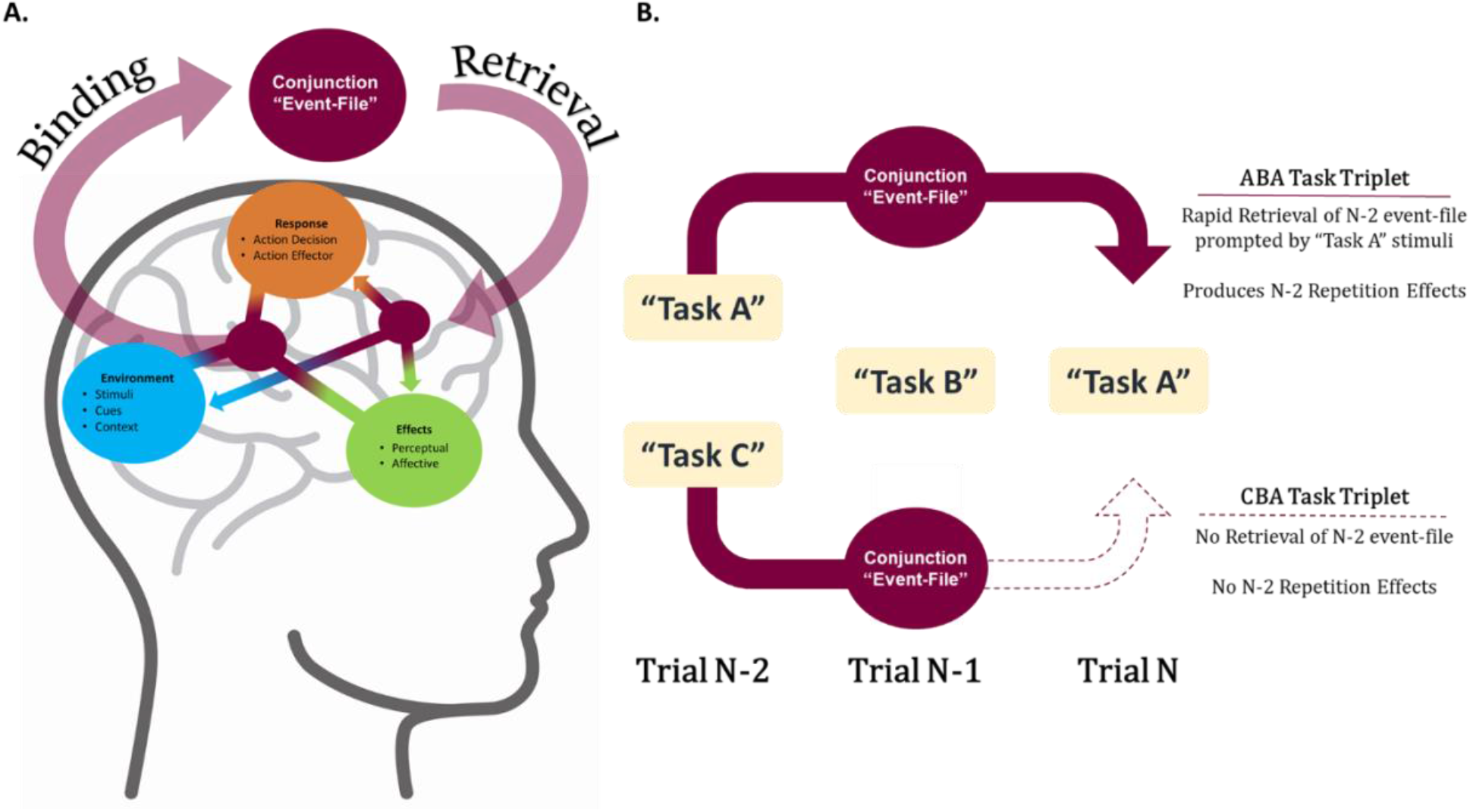
The event-file hypothesis of N-2 repetition costs. A). Diagram of how individual task features are bound as event-files, and then retrieved at a later time. B). Generic task diagram for ABA and CBA trial triplets. Binding of the event-file in trial N-2 occurs for both Tasks A & C, but Retrieval of N-2 event-file only occurs in trial N of an ABA triplet, not a CBA triplet.

However, there is ongoing debate over the source of the N2RC and whether it indeed relates to the formation of event-files (Grange 2018; Grange et al. 2017; Kessler 2018; Kowalczyk and Grange 2020; Scheil and Kleinsorge 2018, 2021). For example, an influential alternative account attributes N2RC effects to the inhibition of task set A by the intermittent switch to task B. This ostensibly impairs the return to the previous, now-inhibited task context (Koch et al. 2010; Mayr 2002; Philipp and Koch 2006; Schuch and Grange 2014; Sexton and Cooper 2017).

In the current study, we address this question from a neuroscientific perspective, using a recently developed method for the quantification of the exact type of conjunctive task feature representation that is predicted by the event-file framework (Kikumoto & Mayr, 2020). Single-neuron recordings and fMRI have already suggested that such conjunctive representations may exist in both non-human (Parthasarathy et al. 2017; Rigotti et al. 2013) and human brains (Kühn et al. 2011). Kikumoto and Mayr’s (2020) method uses multi-variate pattern analysis (MVPA) and representational similarity analysis (RSA) to non-invasively track the emergence of such representations with millisecond precision. It also allows for a direct test of the relationship between the strength of these representations and behavior via linear mixed-effects modeling of trial-to-trial reaction time. Kikumoto and Mayr demonstrated that conjunctive representations decoded from EEG recordings can be used to predict reaction time on the current trial: stronger conjunctions between task context, stimulus information, and response activity accompany faster reaction times.

Here, we leverage this same technique to investigate if conjunctive representations can also account for N2RC effects – i.e., whether the formation of a conjunctive event-file on one trial has adverse effects on future trials, as predicted by the event-file account of the N2RC effect. Using an N2RC paradigm, we first aimed to replicate Kikumoto and Mayr’s finding that conjunctions of task context, stimulus, and response accompany faster reaction times on the same trial. We hypothesized that within ABA triplets, the strength of the conjunctive representation formed on the first instance of task A would be detrimental to reaction time on the second instance of task A if a different response was required. This would provide direct evidence for the N2RC effect being partially attributable to previously formed, now-outdated event-files. More broadly, it would indicate that while event-file formation is typically beneficial to immediate behavior, it can also be detrimental when the behavioral requirements associated with a specific context subsequently change.

## Methods

### Participants

35 healthy young adults participated in the experiment (Age mean(SD) = 19.6 (3.0), 1 left handed, 19 females). Participants were either paid $15 per hour or received course credit for their participation in the study. All participants had normal or corrected-to-normal vision. The experiment was approved by the ethics committee at the University of Iowa (IRB #201511709).

### Materials and procedure

Experimental stimuli were presented via a Ubuntu Linux computer, running Psychtoolbox 3 (Brainard et al. 1997) under MATLAB 2015b (The-MathWorks, Natick, MA). Participants sat upright with their arms and the computer’s keyboard resting on a supportive platform placed on the arm rests of the chair.

### Experimental paradigm

Our paradigm was adapted from Grange et al. (2017, Experiment 2; ***Figure 2***). A large hollow square outline (the frame) remained on the screen across trials. Trials began with the presentation of the task cue (the outline of one of three shapes) inside the center of the frame, followed by a 500ms Cue-Target Interval (CTI). The target (filled black dot) would then appear in one of the four corners of the frame.

**Figure 2.**
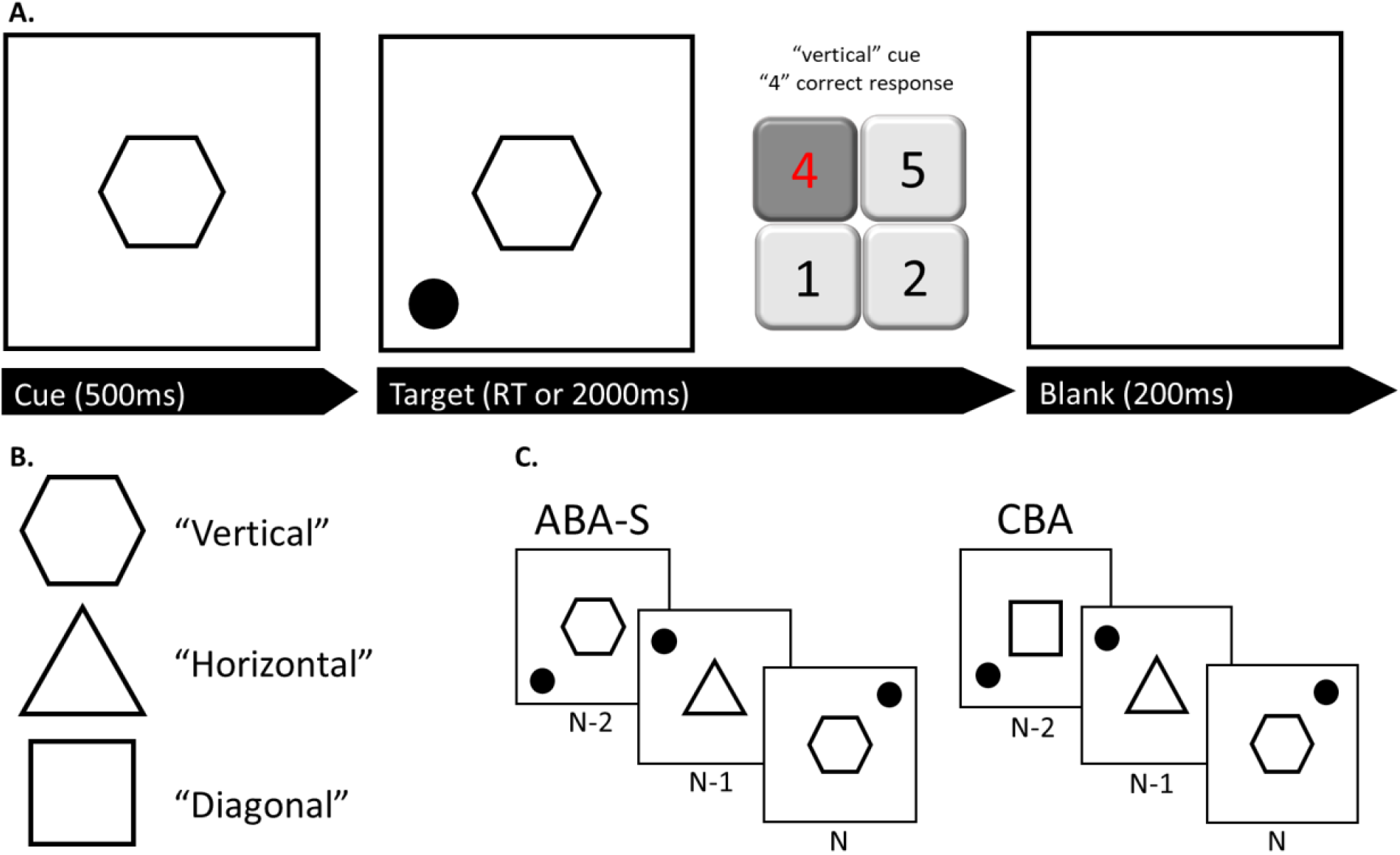
Experimental paradigm. A). Diagram for single trial. The shape cue is presented for 500ms, prior to the target (black dot) onset. The target will be displayed for 2000ms, or until a response is made. Subjects responded using the number pad on a full keyboard. Response is immediately followed by a 200ms Blank ITI before the next cue. B). The three possible shape cues, and their meanings for the task. C). Examples of ABA-S and CBA trial triplets.

Based on the combination of cue shape and target location, participants had 2000ms to respond using their right index finger. The cue shapes “Hexagon”, “Triangle” and “Square” instructed one of three different response mappings (tasks), which corresponded with “Vertical”, “Horizontal” and “Diagonal” movement of the target to one of the unoccupied corners of the frame. For example, should the target appear in the top-left corner of the frame, a triangle-cue would imply that the target should move horizontally to the top-right, hence cueing the response associated with that location. Participants responded using their right index finger and the “4”,”5”,”1” and “2” keys on the keyboard number pad, which corresponded to the top-left, top-right, bottom-left and bottom-right corners of the frame, respectfully. Participants were asked to return their index finger to the center of the response keys after reach trial.

Responses were followed by a 200ms blank period where only the frame remained on screen. If participants failed respond within 2000ms, or made an incorrect response, the usual blank period was replaced by a 500ms presentation of “Faster” or “Error” in red text. The experiment consisted of 800 trials (8 blocks). Cue identity and target location were pseudorandomized to prevent immediate cue (and task) repetitions, and an equal number to task conditions (50% ABA triplets, 50% CBA triplets).

The first two trials of each block were removed from the analysis, as they could not be considered part of a triplet. The behavioral data were then screened for response errors/omissions, RTs lower than 150ms (3 trials overall), and RTs outside 2.5 SDs of the mean (2.33% overall). These trials were removed from further analysis, along with the two subsequent trials. There were three possible task triplet conditions of interest (ABA-S, ABA-R, CBA) with the “-S” and “-R” denoting either a switch or repeat in the required response between trials N-2 and N of an ABA triplet (in principle, CBA-S and CBA-R also exist, but since there is no repetition of task cues within these triplets, the overlap of event files should be zero, regardless of response requirement). Our main comparison of interest was between ABA-S trials and CBA trials. ABA-R trials were of secondary interest, as there was only a limited amount of those triplets per subject (as target locations were drawn from a uniform distribution so as to not introduce a bias, resulting in only one ABA-R triplet for every 3 ABA-S triplets). The dependent variables of interest were RT and accuracy on the final trial of ABA-S and CBA triplets, which were compared using paired-samples t-tests.

### EEG Recording

EEG data were recorded using a 64-channel active electrode cap connected to an actiCHamp amplifier (BrainProducts, Garching, Germany), at a rate of 500 Hz (10 s time-constant high-pass and 1,000 Hz low-pass hardware filters). Electrodes Pz and Fz served as the reference and ground, respectively.

### EEG preprocessing

Raw EEG data were preprocessed utilizing custom MATLAB scripts and EEGLAB toolbox functions (Delorme and Makeig 2004), RRID: SCR_007292. Individual participant data were imported, resampled to 250Hz, and filtered using Hamming windowed sinc FIR filters (pop_eegfiltnew.m; high-pass: 0.01Hz, low-pass: 50Hz). The resulting data were then epoched into two datasets, one target-locked [-600:800ms] and the other response-locked [-700:200ms], which were used to investigate the emergence of task-feature representations relative to either event. Epochs containing non-stereotypical artifacts were then removed from further analysis (joint probability and joint kurtosis; 5.5 SD cutoffs, cf., (Delorme et al. 2007)). Data were re-referenced to a common average prior to ICA decomposition (Infomax; (Bell and Sejnowski 1995)) with extension to sub-gaussian sources (Lee et al. 1999). Components representing eye-movement and electrode artifacts were identified using outlier statistics and removed from the data. Unlike Kikumoto and Mayr (2020), EEG data did not undergo time-frequency analysis, and the following analyses were performed on the raw EEG channel data (i.e., the event-related potential).

### Linear Discriminant Analysis (LDA)

LDA was run on one subject at a time. Each subject had twelve randomly selected trials held out as test data, while their remaining trial data were used for training. Training trial data were then sorted into twelve classes; one for each unique combination of cue, target, and response (i.e., the ‘event file’ or “constellation” in Kikumoto & Mayr, 2020). In the case of unequal numbers of trials across these classes, trials were randomly excluded to achieve equal numbers. The excluded trials were randomly paired with the remaining trials in their respective classes and averaged. This 1-1 pairing and averaging allowed for all data to be included in each training and testing loop, while also ensuring a LDA with balanced classes. LDA was carried out using MATLAB’s *fitdiscr* function. Each training-testing loop applied the LDA to each time-point, with all 63 EEG channels included as features. Posterior probabilities for the hold-out data (12 trials), were saved for each time-point. This process was repeated with a new set of 12 hold-out trials until each trial had been tested. This entire process was then repeated for ten iterations, resulting in each trial being included in the testing set ten times.

### Representational Similarity Analysis (RSA)

The posterior probabilities for all trials and iterations were smoothed using an overlapping moving median window of 20ms. As in Mayr and Kikumoto (2020), the posterior probabilities of each trial for each time-point were regressed onto model vectors representing the idealized classification probabilities for the trial’s true class identity (***Figure 3***). Specifically, four vectors were used as predictors in the regression, representing the cue, target, response, and the conjunction of all three (i.e., the event-file). The t-values of these predictors were saved as measures of similarity. A fifth vector of z-scored/averaged reaction times for each class was also included to account for variations in RT. Following RSA, the median t-value for all predictors at each time-point of each trial was taken across all ten iterations.

**Figure 3.**
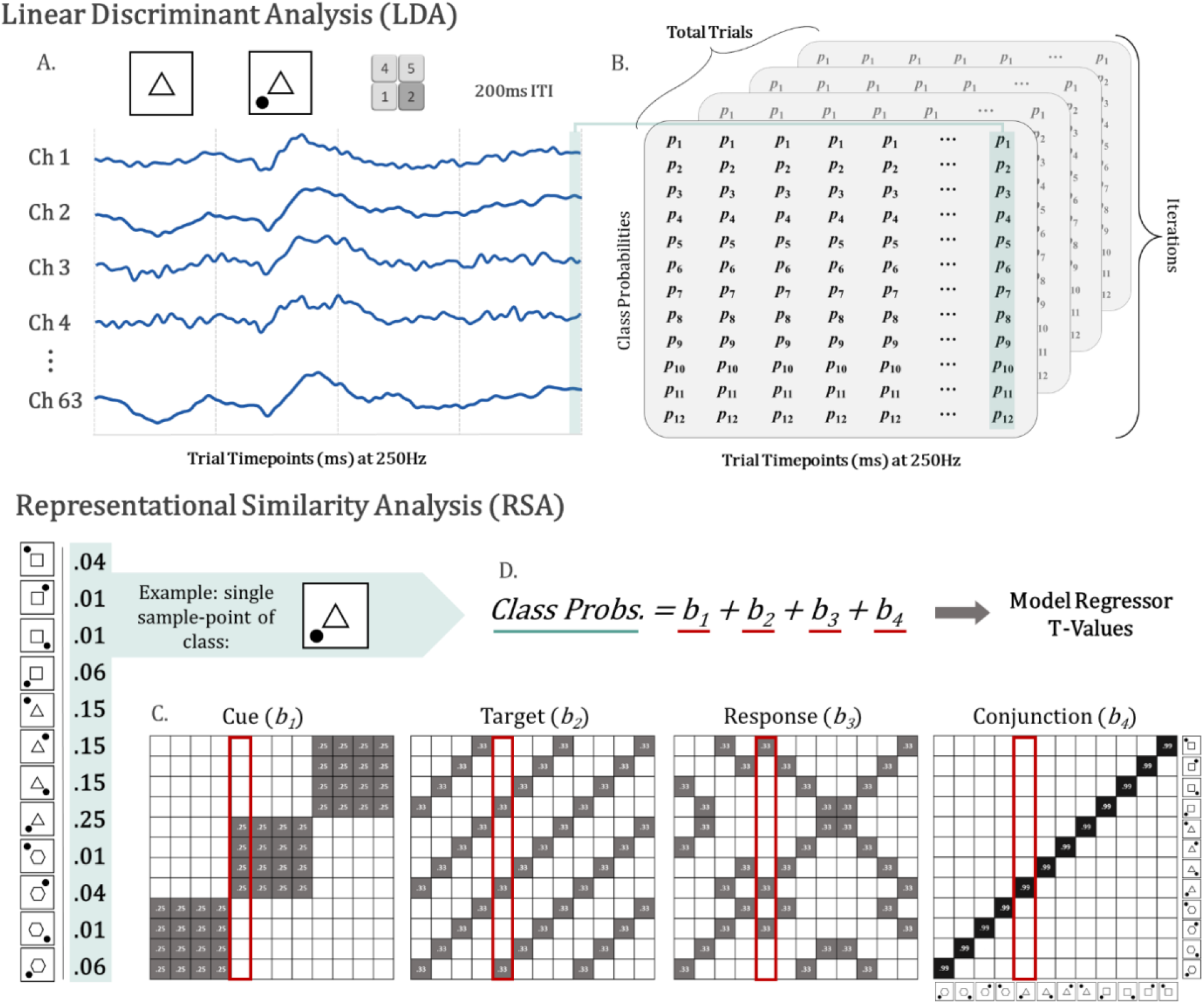
Analysis logic, analogous to Kikumoto & Mayr (2020). A). LDA was run on single subject EEG data, using all 63 channels as features. Each timepoint was decoded (trained/tested) individually, across all trial epochs. B). Posterior probabilities indicating how likely a single sample belonged to each of the 12 classes, were saved in a 4D matrix (Class x Time x Trials x Iterations). C). The RSA consisted of taking individual sample points and regressing them onto the corresponding vector from idealized model matrices. The T-Values for each predictor in the regression D) were saved as a measure of similarity. Once all samples had been regressed, the median T-Values across the 10 Iterations were used for further analysis.

### Mixed-Model analysis

As in Kikumoto and Mayr (2020), the representational strengths of each of the four factors of interest (cue, target, response, conjunction) in the neural data were related to behavior (reaction time) using a mixed effects model. T-values from the RSA were baseline corrected (first 100ms) for each subject. Values outside ± 5*SDs the mean were excluded, and the data was then smoothed using an overlapping 80ms moving mean window. Subject RTs were then log-transformed (ln) and detrended to remove any linear and/or quadratic trends. Subjects’ data were first modeled with the RSA values for cue, target, response, and conjunction added as fixed effects predicting the RT on the same trial. ‘Subject’ was included as the only random effect (***Equation 1***). Subject behavioral data and RSA values were then trimmed to include only trials which made up a complete ABA or CBA triplets (uninterrupted by behavioral or EEG rejections). Two models were created to predict trial N RT, one using trial N-2 data predictors, and the other N-1 predictors. In both cases the N-1 or N-2 RSA values for cue, target, response, and conjunction were added as fixed effects, as well as the trial N condition (ABA-S or CBA). Interactions for all RSA values with the trial N conditions were also included. ‘Subject’ was included as the only random effect (***Equation 2***). This analysis was applied to all time points of the trials. All mixed model analysis were carried out in R (version 4.0.4, packages: *car, lme4*).

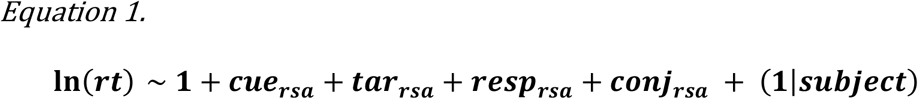

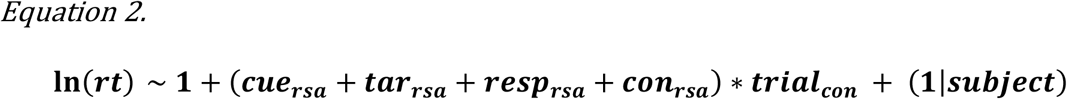

### Cross-Correlation Analyses

To investigate the relationships between the strengths of the neural task representations across the trials of a triplet, the same response-locked data used in the final mixed-model (***Equation 2***) were then cross-correlated for each subject. Trial triplets were sorted into three condition groups (ABA-S, ABA-R, CBA). The RSA values for all task features at every time point in trials N-2 and N-1 of each group, were compared to the corresponding RSA values at every time point in trial N, using Pearson correlations. This produced a 225×225 matrix of Pearson’s r values comparing the task features in trials N-2 and N-1 to those in trial N, for each condition and subject. These matrices were Fisher’ z transformed prior to plotting and further testing (Fisher 1915). The difference between the cross-correlation matrices comparing the conjunctions in trial N-2 and N for ABA-S and CBA triplets, was tested using a one-sided permutation test for dependent measures (10000 permutations, Gerber, 2022). A difference matrix (ABA-S – CBA) was the produced for each subject, and then averaged across both dimensions, post-baseline (first 100ms), resulting in a single value per subject. This was correlated with each subject’s N2RC (Pearson’s).

## Results

### Behavior

Subjects failed to respond on 1.09% of trials and committed response errors on 4.05% of trials overall. As predicted, a comparison of ABA-S and CBA trials reveals N-2 Repetition Cost effects on both RT (ABA-S: mean = 748ms, SE = 26.3; CBA: mean 707.4ms, SE = 26.1) and accuracy (ABA-S: mean = 94.2%, SE = .82, CBA: mean 95.1%, SE = .74). Paired-samples t-tests revealed that these differences were highly significant for both RT (t(34) = 7.48, p < .001) and accuracy (t(34) = −3.74, p < .001).

### Decoding conjunctive representations on the current trial

In a first step, we aimed to replicate the results from Kikumoto and Mayr (2020) to validate that we could identify the conjunctive representation of cue, target, and response (i.e., the event-file) from the neural data. Following EEG epoch rejections, each subject had an average of 517.4 (+/- 58.0) cue/target-locked trial epochs and 515.7 (+/- 58.2) response-locked trial epochs. From the EEG data of each subject, we successfully produced significant point-by-point estimates of representational strength for all task features (cue, target, response) and their unique conjunction (event-file). Single subject RSA averages (T-Values from regression of models onto LDA output) were plotted as a grand-average, time locked to both cue/target onset (***Figure 4a***) and response onset (***Figure 4b***). Notably, the representations for the task cue, target and conjunction were significantly decoded across both time series. The response representation was only significantly decoded when the data was response locked, likely due to variance in RT (***Extended data, Figure 4-1***).

**Figure 4.**
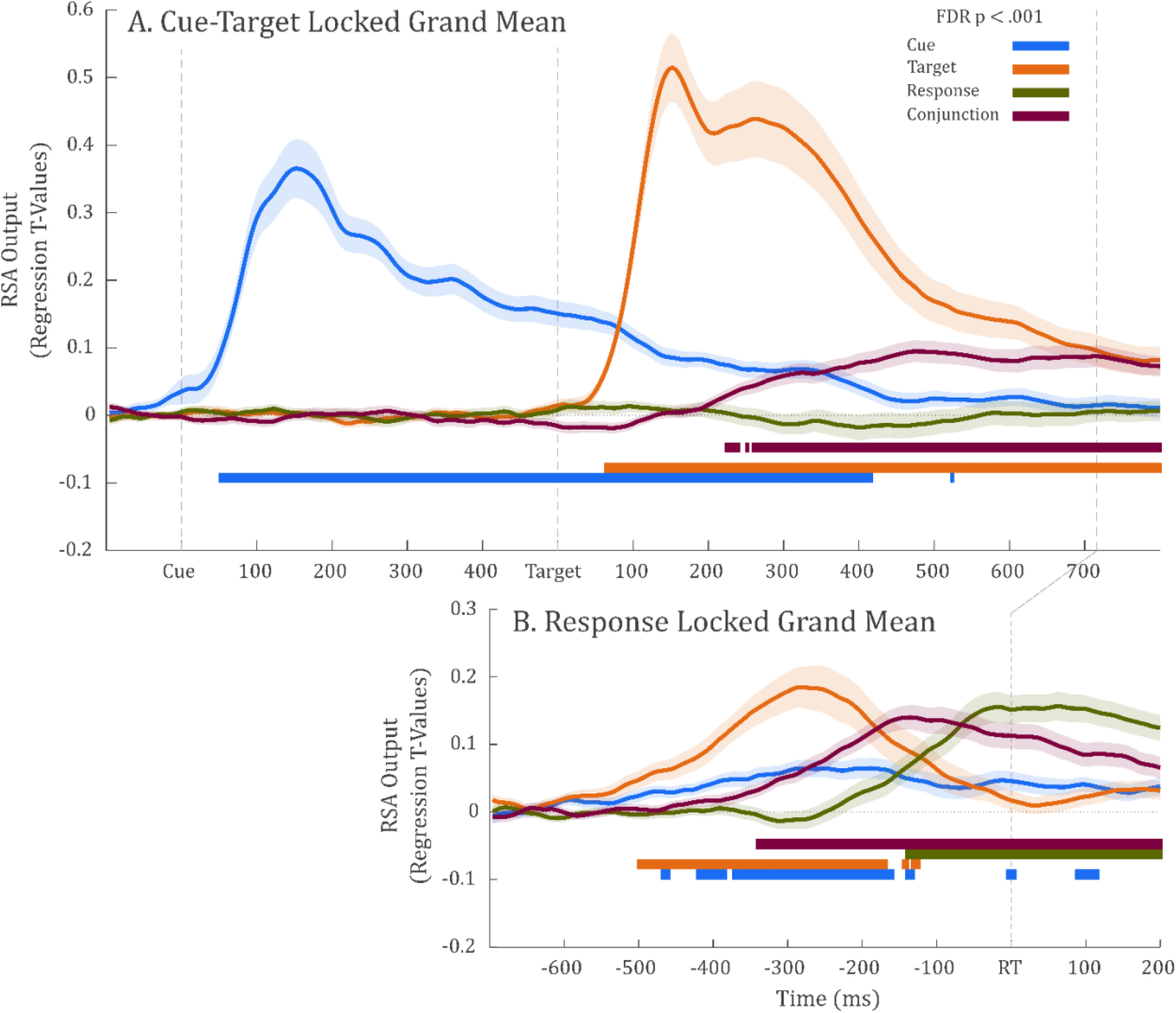
Decoding current trial representations of cue, target, response, and their conjunction from the neural data. A). Grand average of the cue/target locked RSA output. B) Grand average of response-locked RSA output.

Furthermore, the mixed-effect model approach produced results similar to those of Kikumoto and Mayr (2020; ***Figure 5***). Analysis of the response-locked RSA values (***Figure 4b***) showed that increased strength of the cue, target and conjunctive representations predicted faster RTs on the same trial. The response representation did not produce a significant relationship by itself. (However, it did in the cue/target-locked data; see extended data, ***Figure 5-1***).

**Figure 5.**
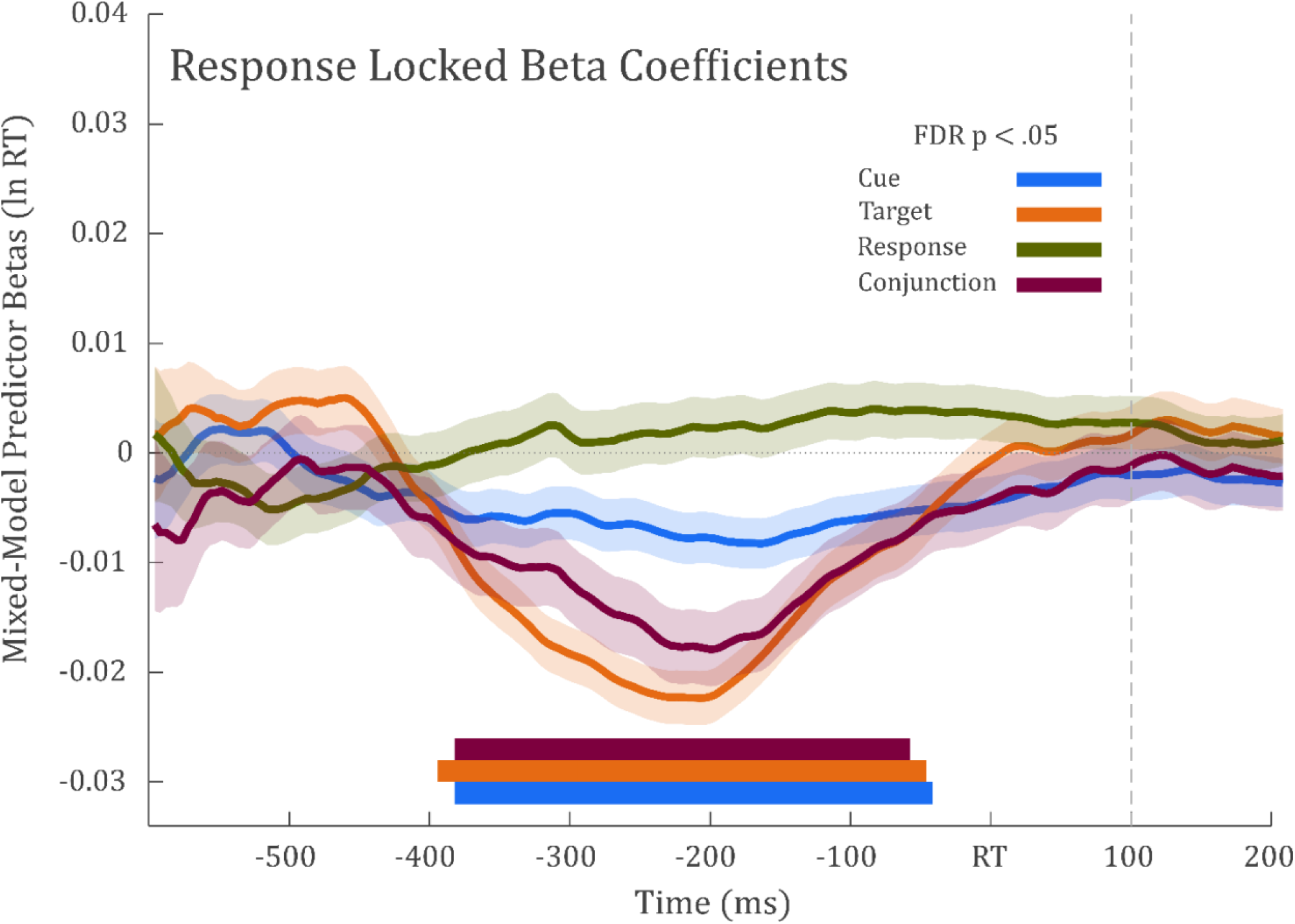
Relationship between behavior and neural representations on the same trial. Plot shows the beta coefficients from the mixed-model analysis of response locked RSA values, fitted at each time point for all subjects and trials. Negative beta values indicate that greater representational strength predicts faster RT during the same trial.

### Main analysis: Trial N-2 conjunctions explain RT cost on trial N

We then tested our main hypothesis – i.e., that in ABA-S triplets, the neural representation of the conjunction (event-file) on trial N-2 would negatively affect the reaction time on the current trial N. A mixed-effects model was used to investigate the relationship between the trial N-2 representational strength of each task factor (cue, target, response, conjunction) and the reaction time of trial N, using the response locked RSA values. The beta coefficients for the interaction terms of each model were plotted at each time point in ***Figure 6***. As hypothesized, the strength of the conjunctive representation in trial N-2 significantly predicted increased trial N RTs. Significant time periods were found from −175 to −83ms and −63 to 33ms relative to the response, for ABA-S triplets compared to CBA triplets (FDR < .05, ***Figure 6, left panel***). In other words, the greater the strength of the conjunctive representation on trial N-2, the slower the RT of trial N in an ABA-S triplet. This was only true for the conjunctive representation, and not for any individual factor (cue, target, response) by itself. Moreover, the conjunctive representation on the immediately preceding trial N-1 did not predict anything about trial N RT (***Figure 6, right panel***). Beyond these interactions, the mixed-model analysis did not show any simple main effects that survived FDR correction (p < .05; ***Extended data, Figure 6-1***).

**Figure 6.**
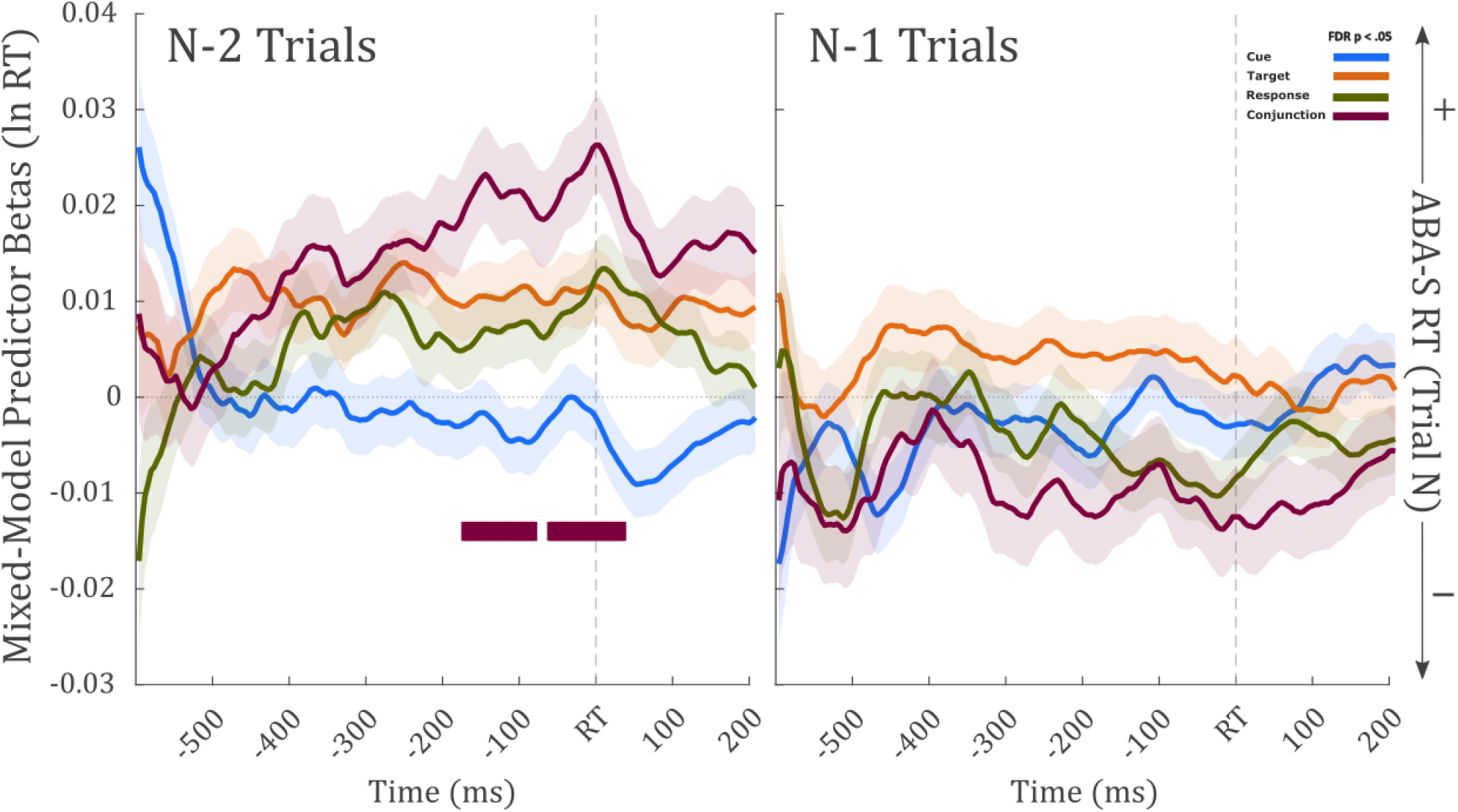
Plots show the beta coefficients of the interaction terms (trial condition x RSA output) from the mixed-model analysis, fitted at each time point, across all subjects and trials. Positive beta values (left y-axis) indicate that increases in the RSA values (greater representational strength) on trial N-2 correspond with slower trial N RT in ABA-S trials, compared to CBA trials (right y-axis). The left plot shows the output of the mixed-model which only included the RSA values from trial N-2 as predictors for trial N RT. The right-plot shows the same, but for the RSA values of N-1 trials only.

### Exploratory analysis: Quantifying cross-trial interactions between conjunctive representations

To further explore the cross-trial dynamics that give rise to the N2RC, we calculated cross-correlations between the representational strengths of the conjunctive representations on the first and third trials within ABA and CBA triplets (***Figure 7***). First, to investigate the critical contrast underlying the N2RC, we compared the correlation of neural conjunctive representation strength between the first and third trial of an ABA-S triplet to the same trial-combination in a CBA triplet. This revealed a marginally significant cluster (***Figure 7A***) that began within ∼100ms before the response. The triplet-specific cross-correlation matrices revealed that this was due to the presence of a consistent anti-correlation in representational strength of the conjunctive representation on CBA triplets, which was absent on ABA-S triplets. In other words, in CBA triplets, stronger conjunctions on trial N-2 corresponded to the formation of weaker conjunctions on trial N, suggesting interference between the different event files corresponding to tasks C and A. This effect was absent on ABA-S trials, whose event files partially overlap. Moreover, these differences in correlation directly related to the size of the N2RC across subjects – i.e., subjects with greater differences in these cross-correlations between CBA and ABA-S trials showed greater N-2 repetition costs (***Figure 7B***). Control analyses revealed that the same was not true for the correlations between non-conjunctive representations (cue, target, response).

**Figure 7.**
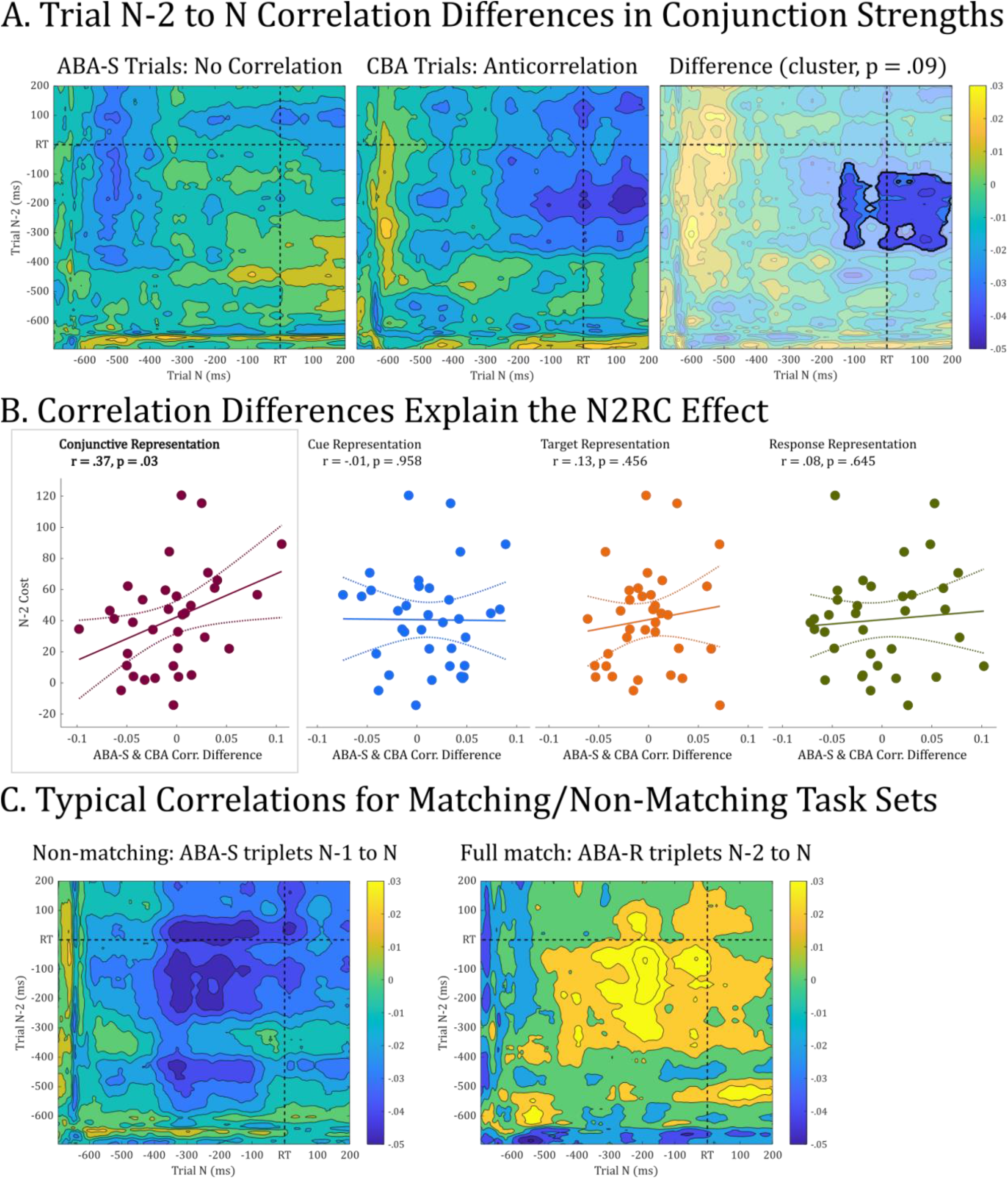
Exploration of the representational cross-correlation matrices. A). The grand mean of the Pearson’s correlation matrices, comparing the conjunctive representation in trials N-2 and N, for both ABA-S and CBA triplets. The difference between these matrices is on the far right, with the highlighted region showing the marginally significant cluster. B). Scatter plots show the relationship between subjects’ averaged difference in ABA-S and CBA cross-correlation matrices (N-2 & N) and their individual N2RC, for all task features. Only the conjunction shows a significant relationship (p = .03). C). These example matrices show the distinct patterns of non-overlapping (ABA-S; N-1 & N) and completely overlapping (ABA-R; N-2 & N) conjunctive representations.

To buttress this analysis, we explored whether the correlation patterns of other trial combinations matched the abovementioned interpretation. First, within ABA-S triplets, the conjunctive representations on trial N-1 and trial N (i.e., non-matching event files) indeed showed the same anti-correlation pattern that was found for N-2 to N combinations within CBA triplets (***Figure 7C, left panel***). This suggests that whenever there is a mismatch in task features as early as the cue (i.e., a task switch), a stronger conjunctive representation of a past task feature combination will impair the formation of the new conjunction on the current trial. Second and further in line with this, ABA-R trials (in which the task sets are identical between trial N-2 and trial N) showed a <u>positive</u> correlation pattern between trials N and N-2 (***Figure 7C, right panel***).

Taken together, these correlation patterns suggest that the formation of a neural representation of the task feature conjunction on a past trial (N-2) benefits the formation of the conjunctive neural representation on the present trial (N) if the task sets match, and is detrimental to the formation of a different conjunction if they mismatch. Moreover, the cross-subject correlation suggests that the strength of these processes is directly proportional to the size of the N2RC effect. In contrast to our main analysis, these exploratory analyses were performed post-hoc and therefore have to be interpreted with caution. However, they generate further insights into the neural dynamics of how the trial N-2 task set may affect behavior on trial N to affect the N2RC.

## Discussion

In the current study, we used a recently established neurophysiological method for tracking conjunctive representations between specific task contexts, stimuli, and responses (i.e., ‘event-files’, Hommel et al., 2001) to address an ongoing debate on the role of such event-files in task switching. Based on the methodological groundwork of Kikumoto and Mayr (2020), we leveraged the combination of RSA and MVPA of EEG data to investigate the trial-to-trial relationship between the strength of conjunctive ‘event-file’ representations and behavior. However, while Kikumoto and Mayr’s investigation was limited to the current trial, we directly tested whether the formation of an event-file on the current trial can influence future behavior – moreover in adverse fashion. Using the N-2 Repetition Cost as a behavioral model, we found that the strength of the conjunctive representation of cue, target, and response on the present trial indeed adversely influences performance on a trial in the near future – specifically, in cases in which response requirements changed between two instances of the same task.

Furthermore, the exploration of correlation patterns between the strength of conjunctive representations on different trials within CBA and ABA triplets generated additional insights into the dynamics by which this adverse influence of past event-files may manifest. First, the comparison of non-matching task contexts (trial N-2 vs. trial N comparisons within a CBA triplet; trial N-1 vs. trial N comparisons within ABA triplets) revealed a stereotypic pattern of decorrelation between the strength of the respective conjunctive representations of successive, different task contexts. In other words, the stronger the representational strength of feature conjunctions on the current trial, the weaker the representational strength of feature conjunctions of subsequent non-matching trials. This suggests that once-formed conjunctive representations may generally interfere with the formation of feature conjunctions on subsequent trials – a phenomenon that may underpin task-switch costs more generally. Moreover, the comparison of task C and task A conjunctions of CBA triplets revealed that this pattern extended beyond the immediately subsequent trial – i.e., stronger representations of feature conjunctions on a given trial still impair the formation of feature conjunctions two trials later. Most interesting, the exploration of ABA-R triplets provided a possible explanation as to why this takes place, as it showed the opposite pattern between the N-2 and trial N conjunctions (i.e., between two spaced trials with matching event-files). In such triplets, stronger conjunctive representations on the first instance of task set A (trial N-2) correlated with stronger conjunctive representations on the second instance (trial N). In line with the general event-file hypothesis, this suggests that the adaptive function of this apparent forward-propagation of a current task set is to benefit behavior if the task set repeats / remains the same. However, if the task set switches, the anti-correlation pattern ensues. Such an automated storage-and-retrieval process makes sense in everyday scenarios the brain has evolved to handle, in which task sets do not typically change with the same regularity as in laboratory experiments like the current one. ABA-S trials then form a middle ground between the mismatch scenarios provided by CBA triplets on the one hand and the full-match scenarios provided by ABA-R trials on the other: the decorrelation observed on mismatch trials does not take place, as the strength of the N-2 conjunction does not adversely influence the formation of the task N conjunction (***Figure 7***). However, as the trial unfolds and the target indicates that response requirements have changed compared to the most recent presentation of the cue, conflict ensues and results in the behavioral cost expressed in the N2RC effect. Indeed, as seen in ***Figure 7B***, the difference in correlation between N-2 to N trial conjunctions in ABA-S triplets compared to CBA triplets (i.e., the main contrast of interest in the current study) was directly correlated with the N2RC effect across subjects. This correlation is very much in line with the results of our main analysis – i.e., that the strength of N-2 conjunctive representations predicted longer reaction times in trial N of an ABA-S triplet. Indeed, subjects for whom the conjunctive trial N-2 representation was stronger – and therefore, interfered more with the formation of the conjunctive trial N representation within a CBA triplet (or primed the response contained in the trial N-2 event-file to a greater extent on trial N within an ABA-S triplet) – showed a larger N2RC effect.

With regards to our main finding –the inverse relationship between the strength of the conjunctive representation on trial N-2 and RT on trial N of an ABA-S triplet – two observations are key to rule out alternative explanations. First, the conjunctive representation of the intervening trial N-1 in an ABA-S triplet did not predict RT on the current trial. This confirms that the N-2 Repetition Cost is indeed specific to a precise match between task contexts. Second, neither the cue nor target representations of trial N-2 by themselves could predict behavior on trial N. This shows that regardless of how strongly the target or cue was encoded on trial N-2, only the strength of their binding with the task context in a conjunctive representation affected future behavior. As such, our results are consistent with previous work, suggesting that the episodic retrieval event-files significantly contributes to the N2RC effect (Frings et al. 2020; Grange 2018; Grange et al. 2017; Hommel 1998; Kühn et al. 2011; Neill 1997; Spapé and Hommel 2010). Additionally, we are the first to demonstrate that a direct measurement of task representational strength from neural data can be used predict future RT costs.

As such, we successfully used the decoding of conjunctive representations from neural recordings to not only show that greater task-feature binding can both be beneficial to current behavior and detrimental to future behavior, but we also demonstrated how neurophysiological methods can provide crucial insights into behavioral phenomena that have hitherto mainly been investigated using experimental psychology. Moreover, our results provide a powerful demonstration of the event-file framework in explaining behavior and linking it to neural activity. Indeed, conjunctive representations of stimuli, actions, and outcomes appears to be a fundamental code that the human brain uses to guide future behavior. While early conceptualizations of event or ‘object’-files focused on the conjunction of visual stimulus representations (Kahneman et al. 1992; Treisman 1998), the idea that actions and their outcomes can be commonly linked together with perceptual information is hence a key insight into human adaptive behavior (Hommel 1998; Hommel et al. 2001; Spapé and Hommel 2010). This proposal sheds light on how event-files allow us to do more than simply recognize a familiar sensory stimulus, but also contextualize that stimulus by its expected effects, our own actions, and the surrounding environment.

One open question that remains is the role of backwards inhibition in the N2RC effect. Past studies have shown that ABA triplets can show N-2 Repetition Costs even when the current response requirement matches the past response requirement (Grange 2018; Grange et al. 2017), which is a counterintuitive finding under the event-file framework. The magnitude of these costs tends to be smaller than those in ABA-S trials, suggesting that they may represent a “pure” form of backward inhibition which can only be properly examined when the effects of episodic retrieval are controlled for (Grange 2018; Kowalczyk and Grange 2020). While ABA-R trials were not the primary interest of the current study (as the experiment only contained a limited number of ABA-R trials and used a CTI that is too long to expect substantial RT effects on these trials), the framework presented here could be adapted to specifically test this hypothesis. Indeed, neurophysiological signatures of inhibitory processes are well-established in both the motor and cognitive domains (Anderson et al. 2016; Wessel 2019; Wessel and Aron 2017), and it has long been proposed that task switch cues activate inhibitory networks (Jamadar et al. 2010; Koch et al. 2010). The combination of the current approach with direct measurements of inhibition could provide insight into whether there is indeed an additional role for inhibition in the N2RC effect. Moreover, the analytical method developed by Kikumoto and Mayr (2020) could be adapted to identify signatures of the latent conjunctive representations established in trial N-2 before its retrieval on trial N (e.g.,(Rose et al. 2016)) and to investigate whether task switching cues on the intervening trial N-1 directly affect the strength of the latent conjunctive representation.

In summary, we here used the recently developed methodological approach for the measurement of conjunctive representations of task context, stimuli, and responses (Kikumoto & Mayr, 2020) to demonstrate that such representations, once formed, can adversely affect future behavior – here, specifically in the form of the N2RC effect. This not only highlights the strength of this analytical approach, but it contributes to the extant literature on the role of episodic retrieval in N-2 Repetition Costs. Hence, this work provides fundamentally novel insights into a long-standing debate in the behavioral literature and illustrates the potential of this neuroscientific approach to tackling questions of adaptive and flexible behavior.

## Acknowledgements

This work was funded by the National Institutes of Health (NIH R01 NS102201 to JRW).

